# A biomechanical switch promotes lysosomal remodelling and exocytosis in keloid fibroblasts

**DOI:** 10.1101/2023.11.01.564877

**Authors:** Rosie Ross, Yiyang Guo, Rebecca N. Walker, Daniele Bergamaschi, Tanya J. Shaw, John T. Connelly

## Abstract

Keloids are a severe form of scarring for which the underlying mechanisms are poorly understood, and treatment options are limited or inconsistent. While biomechanical forces are potential drivers of keloid scarring, the direct cellular responses to mechanical cues have yet to be defined. The aim of this study was to examine the distinct responses of normal dermal fibroblasts (NDFs) and keloid-derived fibroblasts (KDFs) to changes in extracellular matrix (ECM) stiffness. When cultured on hydrogels mimicking the elasticity of normal or scarred skin, KDFs displayed greater stiffness-dependent increases in cell spreading, F-actin stress fibre formation, and focal adhesion assembly. Elevated acto-myosin contractility in KDFs disrupted the normal mechanical regulation of ECM remodelling, leading to constitutive collagen and fibronectin deposition. Transcriptional profiling identified mechanically-regulated pathways in NDFs and KDFs, including the actin cytoskeleton, Hippo signalling, and autophagy. Further analysis of the autophagy pathway revealed that autophagic flux was intact in both fibroblast populations and depended on acto-myosin contractility. However, KDFs displayed marked changes in lysosome organisation and an increase in lysosomal exocytosis, which was mediated by acto-myosin contractility. Together, these findings demonstrate that KDFs possess an intrinsic increase in cytoskeletal tension, which heightens the response to ECM mechanics and promotes lysosomal exocytosis.

## Introduction

Keloids are an aggressive type of scarring in which the fibrotic tissue overgrows the boundaries of the initial wound and does not regress spontaneously. Keloid scars are induced by cutaneous trauma, such as cuts, burns, acne, chickenpox, vaccinations, or piercings, and they can be itchy, painful, cause cosmetic disfigurement and physical impairment (Andrews et al. 2016a; Robles and Berg 2007; Trace et al. 2016). As a result, keloid patients often report a decrease in both the psychological and physical domains of quality of life (Bock et al. 2006; Furtado et al. 2009).

Current management options, including steroid injections, surgery, radiation therapy, compression therapy and cryotherapy, are broad and often ineffective with high rates of recurrence (Ekstein et al. 2021). Thus, greater mechanistic insight into the pathogenesis of keloid scar formation is needed in order to develop targeted and more effective therapies.

While the underlying causes of keloid scar formation are poorly understood, they likely involve both genetic and environmental factors (Andrews et al. 2016b; Nakashima et al. 2010). Keloids tend to run in families and are more prevalent in Asian and Afro-Caribbean populations (Alhady and Sivanantharajah 1969; Andrews et al. 2016b), and numerous growth factors and inflammatory cytokines have been implicated in scar formation (Ghazizadeh et al. 2007; Kikuchi et al. 1995; Younai et al. 1994). The resulting scar tissue in keloids are characterised by excess deposition of extracellular matrix (ECM) with rigid, densely packed and hyalinised collagen bundles (Lee et al. 2004), alongside a distinct cartilage-like ECM composition with a decrease in proteases compared to normal skin and normal scars (Barallobre-Barreiro et al. 2019).

Mechanical input is essential for normal wound closure, and several studies have shown increased scar formation or pathological scar formation with additional mechanical stress (Aarabi et al. 2007; Chin et al. 2010; Pietramaggiori et al. 2007). Moreover, the well-studied mechanically-induced transcription factors YAP/TAZ, play a key role in scar formation, and YAP/TAZ inhibition promotes regenerative wound healing including recovery of skin appendages, matrix ultrastructure and mechanical strength in mouse models (Mascharak et al. 2021).

Consistent with these findings, keloid scars are more likely to occur at locations of the body that are under high skin tension, such as the chest, the shoulder and scapular regions and the supra-pubic region (Ogawa et al. 2012; Ogawa et al. 2003; Omori et al. 2009). In addition, keloid scars exhibit higher tension at the scar edges, which are often the areas of high inflammation, and keloid scars appear to spread in the direction of skin tension dictating the shape at specific locations on the body (Akaishi et al. 2010; Akaishi et al. 2008). Keloid tissue and keloid dermal fibroblasts (KDFs) are stiffer than normal skin tissue and normal dermal fibroblasts (NDFs), an observation that is lost in KDFs when treated with the myosin II inhibitor, Blebbistatin (Deng et al. 2021; Hsu et al. 2018; Kenny 2016). Mechanistically, the mechano-sensors YAP/TAZ are upregulated and more localised within the nucleus in KDFs compared to NDFs (Aramaki-Hattori et al. 2017; Deng et al. 2021; Gao et al. 2021), and inhibition of YAP/TAZ significantly reduces KDF proliferation, migration and the production of collagen type 1 α-chain 1 (COL1A1) but induces cell apoptosis (Gao et al. 2021).

Collectively, these observations suggest that mechanical stimuli and mechanotransduction play a central role in keloid scar pathogenesis; however, the direct influences of extracellular mechanical cues on keloid fibroblast function are incompletely understood. Therefore, this study investigated the distinct responses of NDFs and KDFs to defined mechanical environments using the well-established polyacrylamide (PA) hydrogel system to tune the stiffness of the ECM. We observed that KDFs were more sensitive to ECM mechanics with greater increases of F-actin stress fibres and focal adhesions on stiff substrates compared to NDFs. Transcriptional profiling also revealed key mechano-sensitive signalling pathways that were altered in KDFs, and through this analysis we identified a switch in lysosomal remodelling and exocytosis, which was biomechanically regulated in the KDFs. Together, these findings provide new insight into the altered mechano-sensing behaviour of KDFs and their potential role in scar pathogenesis.

## Results

### Keloid fibroblasts display enhanced adhesive and cytoskeletal responses to increased ECM stiffness

To investigate the direct influences of ECM mechanics on normal and keloid fibroblast function we employed PA hydrogels as model substrates for tuning the elastic moduli within physiologic ranges. We generated collagen-coated PA hydrogel surfaces with moduli of 8kPa, representing the approximate stiffness of normal skin, or 70 kPa and 214 kPa, representing the stiffness of keloid scars (Kenny, 2016). NDFs from the skin of three normal donors and KDFs from the scars of four keloid patients were cultured on each of these substrates and examined for morphological changes and cytoskeletal remodelling. Whole cell staining with a fluorescent membrane dye indicated that the KDFs increased their cell area and became less elongated on the stiffer substrates, while the NDFs did not display significant changes in cell morphology across the same range of moduli (Fig 1A-C). Further immunofluorescence staining of key intracellular mechanical structures, including the F-actin cytoskeleton and focal adhesions (FAs), revealed overall more prominent F-actin stress fibres and FAs in KDFs compared to NDFs (Figure 1D-E), and quantitative image analysis confirmed a greater stiffness-dependent upregulation in the number of F-actin filaments and FAs in KDFs compared to NDFs (Figure 1D-G). Collectively, these results indicate a stronger mechanical response to increased ECM stiffness in KDFs compared to NDFs.

**Figure 1:**
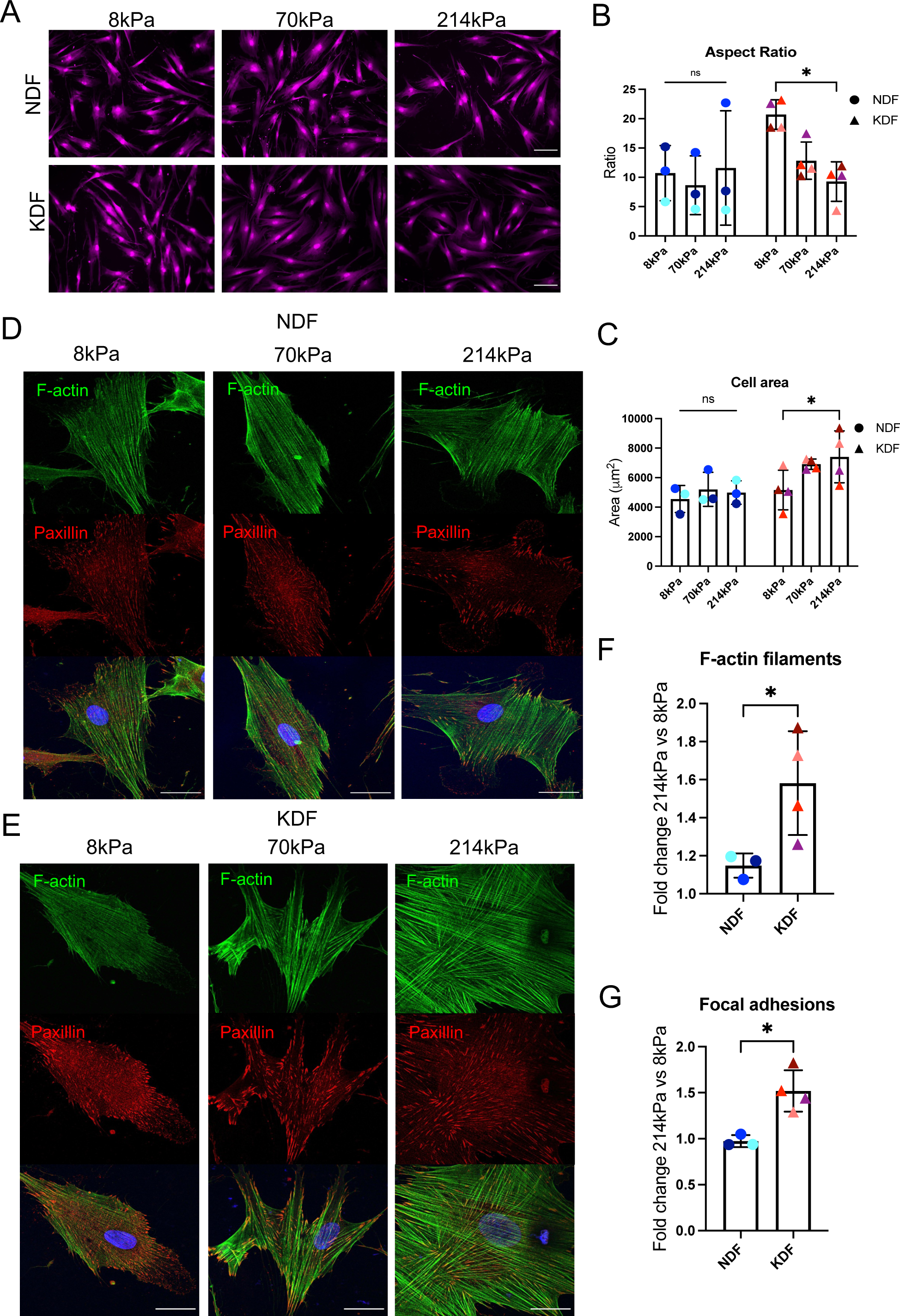
ECM stiffness differentially regulates adhesion and spreading of normal and keloid fibroblasts. Fibroblasts from four different keloid patients (KDF) and three different normal donors (NDF) were grown on soft (8kPa) and stiff (70-214kPa) collagen-coated PA hydrogels for 24 h. (A) Representative images of cells grown on PA hydrogels and stained for cell mask. Scale bar = 100 μm. (B) Quantification of the mean aspect ratio (major:minor axis) of cells. (C) Quantification of the mean cell area. (D) Representative confocal images of paxillin and F-actin (phalloidin) in NDF and (E) KDF. Scale bar = 30 μm. (F) Quantification of the average fold change of the number of F-actin filaments and (G) focal adhesions per cell grown on 214 kPa versus 8 kPa PA hydrogels. Data points represent the mean of individual donors, and bars indicate the overall mean for NDF and KDF cells ±SD. *P<0.05, Two-way ANOVA for B/C and unpaired T-test for F/G.

### Keloid fibroblasts are refractory to ROCK inhibition of ECM deposition

To test the functional implications of altered cytoskeletal remodelling in KDFs, cells were cultured on rigid collagen-coated coverslips and treated with the ROCK inhibitor, Y27632, to disrupt actin polymerisation and acto-myosin contractility. Deposition of cell-derived ECM proteins, type I collagen and fibronectin, was then analysed after five days by decellularisation and immunofluorescence imaging. Here, we observed that treatment with 10 μM ROCK inhibitor completely disrupted F-actin filaments in the NDFs, but not in the KDFs in which large stress fibres were still present 24 hours after inhibitor treatment (Figure 2A). These results are consistent with the heightened response to ECM stiffness and suggest that KDFs possess higher constitutive levels of acto-myosin contractility.

**Figure 2:**
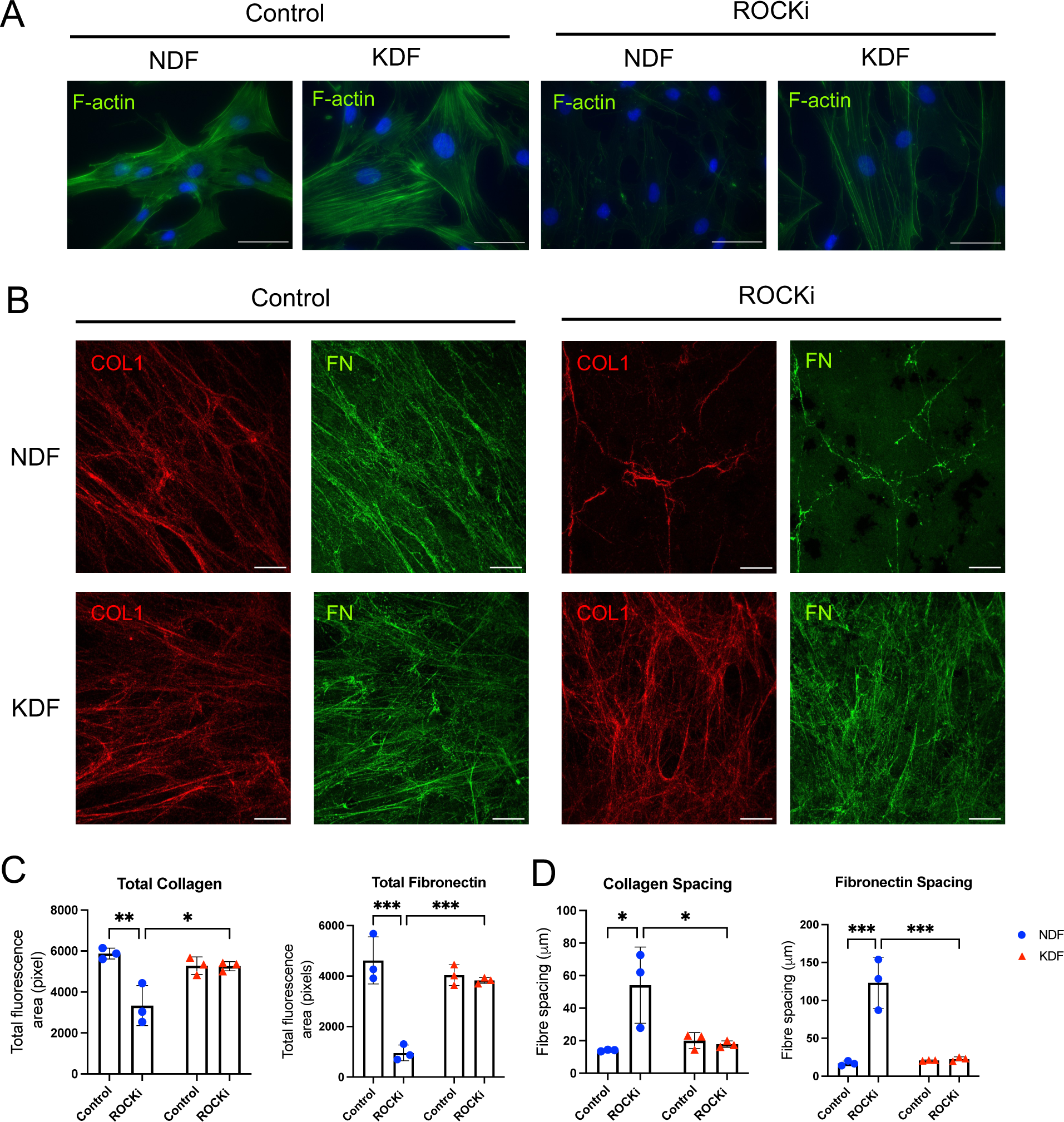
Elevated acto-myosin contractility and ECM deposition by keloid fibroblasts are resistant to ROCK inhibition. (A) NDFs and KDFs were cultured on collagen-coated coverslips for five days in the presence with or without ROCKi (10 μM Y27632). Representative fluorescence images of phalloidin show the F-actin cytoskeleton. Scale bar = 50 μm. (B) Representative confocal images of cell-derived matrices of type I collagen (COL1) and fibronectin (FN). Scale bar = 20 μm. (C) Quantification of total collagen and fibronectin area (pixels) per image. (D) Quantification of mean spacing between collagen and fibronectin fibres (μm). Data represent the mean ± SD from N=3 independent experiments with the representative NDF and KDF donors, N1 and K1. *P<0.05, Two-way ANOVA.

The resistance to ROCK inhibition in KDFs also affected the amount and organisation of cell-derived ECM. In NDFs, treatment with the ROCK inhibitor significantly reduced the total amount of collagen and fibronectin deposited on the surface, and the ECM fibres were more widely spaced compared to untreated controls (Figure 2B-D). By contrast, there were no significant effects of the ROCK inhibitor on collagen and fibronectin deposition by KDFs (Figure 2B-D). Together, these results indicate that acto-myosin contractility facilitates new ECM deposition by dermal fibroblasts, and increased contractility in KDFs confers resistance to ROCK inhibition.

### Transcriptional profiling reveals biomechanically regulated pathways in keloid fibroblasts

Next, we sought to identify downstream mechano-sensitive signalling pathways that were affected by the altered biomechanical behaviour of KDFs via transcriptomic profiling. Both NDFs and KDFs were cultured on soft (8 kPa) and stiff (214 kPa) PA hydrogels and analysed by next-generation RNA-sequencing (RNA-seq). Differential expression analysis was performed between the four experimental groups: NDF 8 kPa, NDF 214 kPa, KDF 8 kPa and KDF 214 kPa, and 539 differentially expressed genes (DEGs) were identified. Principle components analysis (PCA) for the entire gene set was also used to investigate the relatedness between the samples and revealed that the NDFs clustered closely together and away from KDFs along PC1 (Figure 3A). However, KDF samples segregated primarily by patient along PC2 with little visible effect of ECM stiffness, suggesting that high levels of inter-patient variability contribute significantly to KDF transcriptional profiles. To identify mechanically regulated genes, we therefore explored additional principal components just within the DEGs, and by comparing PC5 vs. PC1, we observed that each cell type segregated based on ECM stiffness, which accounted for approximately 2% of the variance (Figure 3B). The DEGs were then filtered to identify the genes that contributed to the variation seen along PC5 (absolute rotation value ≥10, [308 genes]; Supplementary Data File1).

**Figure 3:**
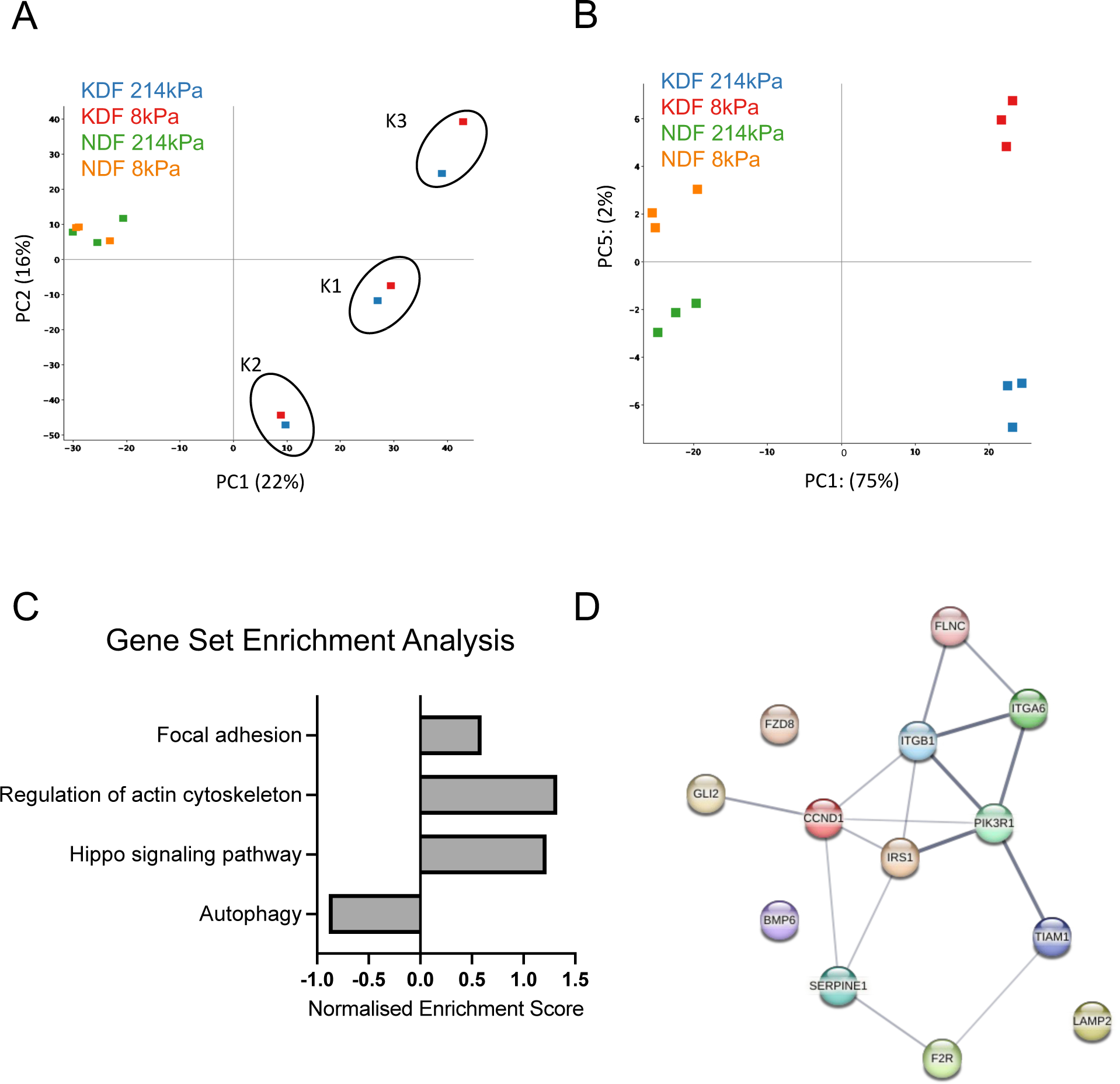
Transcriptomic analysis of mechanically-regulated genes in keloid fibroblasts. Fibroblasts from three keloid patients and three normal donors were cultured on soft (8 kPa) and stiff (214 kPa) collagen-coated PA hydrogels for 24 hours and analysed by next-generation RNA-sequencing (RNA-seq). (A) Principal components analysis (PCA) for all genes showing PC1 and PC2 of each sample. Circles denote keloid cells from patients K1, K2, and K3. (B) PCA for just DEGs showing PC5 versus PC1 of each sample. (C) Gene set enrichment analysis (GSEA) of the highest weighted DEGs (absolute rotation value >10) contributing to PC5, performed by WebGestalt using the KEGG database (top 20 based on FDR adjusted p-value). (D) String analysis of interactions within the enriched genes from the pathways shown from the GSEA.

Gene set enrichment analysis (GSEA) for the highest weighted genes from PC5 was conducted using the Log fold-change in expression in KDFs compared to NDFs and revealed positive enrichment for classic mechano-sensitive pathways, including focal adhesion, regulation of the actin cytoskeleton, and the Hippo signalling pathway (Dupont et al. 2011; Iskratsch et al. 2014) (Figure 3C). Enrichment of genes associated with these known mechanotransduction mechanisms was consistent with the observed differences in cytoskeletal remodelling between NDFs and KDFs and supported the validity of this analysis. Among several positively and negatively regulated pathways (Figure S1), autophagy was a notable classification that was overall downregulated in KDFs compared to NDFs (Figure 3C). Further interrogation of the interactions between the mechanotransduction and autophagy-related genes using String analysis revealed a network with central linkages between genes involved in autophagy (*IRS1, PIK3R1*) and integrin-mediated adhesion (*ITGB1, ITGA6, TIAM1*) (Figure 3D). Taken together, this transcriptional analysis demonstrates that although there is a high degree of heterogeneity within the KDFs, there is still a common mechanically regulated transcriptional response, and autophagy may be an important mechano-sensitive pathway in keloid pathogenesis.

### Autophagic flux is regulated by acto-myosin contractility in normal and keloid fibroblasts

Autophagy, also known as macroautophagy, is a fundamental intracellular process by which defective organelles or misfolded proteins are targeted for degradation by entrapment within the autophagosome, followed by fusion with the lysosome (Jeong et al. 2020). It provides an important mechanism for recycling of biomolecules in response to cellular stress, and several recent studies have implicated autophagy in wound healing and scar formation (Lee et al. 2020; Okuno et al. 2018; Ren et al. 2022). In addition, mechano-sensitive YAP/TAZ signalling promotes autophagic flux via regulation of autophagosome and lysosome fusion (Totaro et al. 2019). Based on our transcriptomic analysis and these previous studies, we therefore hypothesised that altered mechano-sensing may disrupt the normal process of autophagy in KDFs.

To explore the biomechanical regulation of autophagy, NDFs and KDFs were cultured on soft (8 kPa) and stiff (214 kPa) PA gels, and the expression of LC3-I/II was examined by western blotting, as the conversion of LC3-I to LC3-II is a key regulatory step in maturation of the autophagosome. Across all NDFs and KDFs derived from three different donors, there was an increase in LC3-II expression on stiff compared to soft gels, but no significant difference between NDFs and KDFs (Figure 4A-B). However, autophagosome formation is a dynamic process, and increased LC3-II may reflect either an increase in autophagy or a block in autophagosome degradation (Mizushima and Yoshimori 2007). Therefore, we next examined autophagic flux on rigid coverslips using Rapamycin to stimulate autophagy, via inhibition of mTOR, and Bafilomycin to block degradation, via inhibition of autophagosome fusion with the lysosome. Alongside these inhibitors we used the strong myosin inhibitor, Blebbistatin, which in contrast to the previous ROCK inhibitor treatments completely blocked F-actin stress fibre formation (Figure 5C). In both NDFs and KDFs, treatment with Rapamycin did not significantly affect LC3-II expression, and Bafilomycin significantly increased LC3-II (Figure 4C-E), indicating that autophagic flux is active in both cell types. The response to Bafilomycin was significantly dampened by treatment with Blebbistatin in the KDFs, and to a lesser, non-significant degree in the NDFs (Figure 4C-E).

**Figure 4:**
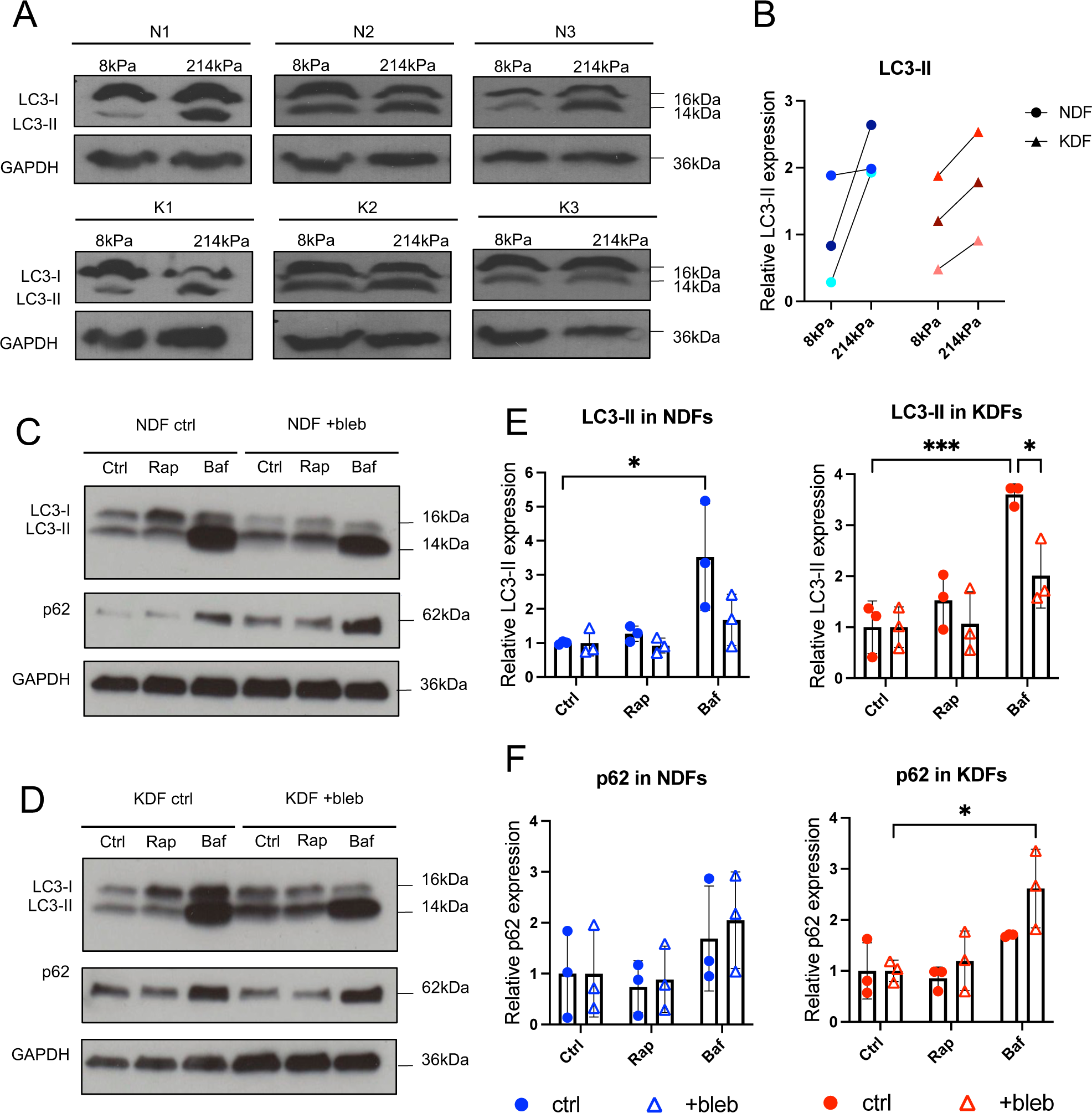
Acto-myosin contractility enables autophagic flux in normal and keloid fibroblasts. NDFs and KDFs (n=3 donors) were cultured on soft (8 kPa) and stiff (214 kPa) collagen-coated PA hydrogels for 24 hours. (A) Western blot analysis of LC3-I, LC3-II and GAPDH in both NDFs and KDFs. (B) Densitometry analysis of LC3-II expression relative to GAPDH. Data points represent values for individual donors, with relative changes between soft and stiff gels linked by a line. (C) NDFs and KDFs were cultured on collagen-coated tissue culture plastic and treated with either DMSO or the myosin-II inhibitor, Blebbistatin (50 μM) for 24 hours. The cells were also subjected to Rapamycin (50 nM) and Bafilomycin A1 (100 nM) treatment for 24 and 4 hours, respectively. Representative western blots show expression of LC3-I, LC3-II, p62, and GAPDH in NDFs and (D) KDFs for control (0.1% DMSO) or inhibitor treatments. (E) Densitometry analysis of LC3-II and (F) p62 expression relative to GAPDH and normalised to the respective control. Data represent the mean ± SD of N=3 independent experiments using donor lines N1 and K1. *P<0.05, ***P<0.0005, Two-way ANOVA.

**Figure 5:**
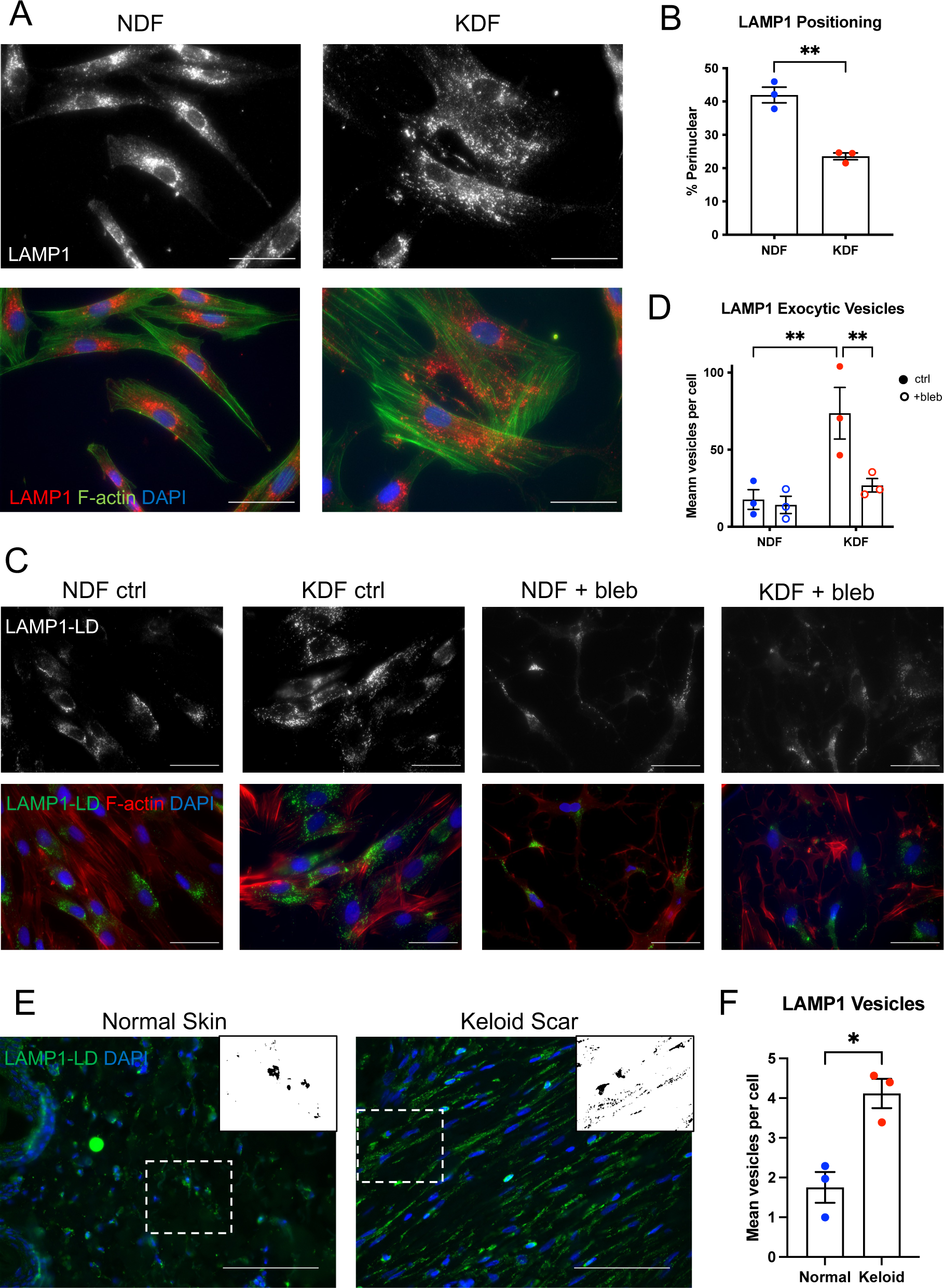
Upregulation of acto-myosin contractility promotes lysosomal exocytosis in keloid fibroblasts. (A) Representative fluorescence images of LAMP1 and F-actin (phalloidin) in NDFs and KDFs cultured on collagen-coated coverslips. Scale bar = 50 μm. (B) Quantification of percentage of LAMP1 within the perinuclear region, defined as the area 50% greater than the nucleus. Data represent the mean ± SEM of N=3 independent experiments using donor lines N1 and K1. *P<0.005, unpaired T-test. (C) Representative images of LAMP1 luminal domain (LAMP-LD) and F-actin (phalloidin) in NDFs and KDFs cultured on collagen-coated coverslips under control (0.1% DMSO) or Blebbistatin (50 μM) treated conditions. Scale bar = 50 μm (D) Quantification of the mean number of exocytic vesicles (LAMP1-LD positive) per cell. Data represent the mean ± SEM of N=3 independent experiments using donor lines N1 and K1. *P<0.005, Two-way ANOVA. (E) Representative images of LAMP1-LD staining of the dermis in normal skin and keloid scars. Inset shows threshold image of LAMP vesicles in region of interest. Scale bar = 100 μm. (F) Quantification of the mean number of LAMP1-LD vesicles per cell in normal skin and keloid scars. Data represent the mean ± SEM of three normal skin samples (N1, N2, N3) and three keloid samples (K1, K5, K6).

Here, we also examined the expression of p62, which is a receptor within the autophagosome that targets proteins for degradation. Activation of autophagic flux will lead to degradation of p62, but a block in the process will result in accumulation (Jiang and Mizushima 2015). Consistent with the effects on LC3-II, Rapamycin had little effect on p62 in both cell types, while Bafilomycin increased p62 expression (Figure 4C-D,F). The level of p62 was further elevated by combined treatment with Bafilomycin and Blebbistatin in the KDFs (Figure 4F). These results provide additional evidence that there is active autophagic flux in both NDFs and KDFs, and that KDFs are slightly more sensitive to inhibition of acto-myosin contractility. Overall, we conclude that ECM stiffness and cytoskeletal tension facilitate autophagic flux in a similar manner for both NDFs and KDFs.

### Lysosomal remodelling and increased exocytosis in keloid fibroblasts depends on acto-myosin contractility

In addition to autophagosome dynamics, we investigated potential downstream changes in lysosome organisation by immunofluorescence staining for lysosomal markers LAMP1 and LAMP2 in NDFs and KDFs. We observed striking differences in the localisation of LAMP1, which localised to the perinuclear region in NDFs but was distributed throughout the cytoplasm in KDFs (Figure 5A-B). Additionally, there was an overall reduction in LAMP2 expression in the KDFs (Figure S2A), consistent with the RNA-seq analysis. Recent studies have shown that peripheral LAMP1 vesicles are associated with lysosomal exocytosis, and exocytic vesicles can be detected by the presence of the LAMP1 luminal domain (LAMP1-LD) on the cell surface (Andrews 2017; Buratta et al. 2020). We therefore stained non-permeabilised NDFs and KDFs with a monoclonal antibody for LAMP1-LD, which labelled clear vesicular structures on the cell membrane (Figure S2B). Under control conditions, KDFs displayed significantly higher numbers of exocytic vesicles compared to NDFs, and disruption of acto-myosin contractility with Blebbistatin significantly reduced the number of vesicles to the level of NDFs (Figure 5C-D). Further analysis of human tissue samples by immunofluorescence staining revealed greater levels of LAMP1-LD vesicles in highly aligned fibroblasts of keloid scars compared to fibroblasts in the papillary dermis of normal skin (Figure 5E-F), suggesting that lysosomal function is similarly altered in vivo. Together, these findings indicate that elevated cytoskeletal tension in KDFs controls lysosomal organisation and promotes a switch to greater exocytic activity.

## Discussion

In this study, we characterised the direct effects of altered ECM stiffness on the cellular responses of NDFs and KDFs. Compared to NDFs, KDFs displayed an intrinsically greater biomechanical response to rigid ECM substrates as demonstrated by increased spreading, formation of F-actin stress fibres, and focal adhesion assembly, suggesting that they possess increased acto-myosin contractility and tension within the actin cytoskeleton. Higher cytoskeletal tension also rendered KDFs refractory to ROCK inhibition and modulation of ECM deposition. Through RNA-seq transcriptional profiling, we identified a core set of approximately 300 mechanically regulated genes that were differentially expressed between NDFs and KDFs, and these DEGs were enriched for a number of distinct molecular pathways, of which autophagy was a notable example. Further analysis of autophagy and the lysosome confirmed the functional importance of this pathway and demonstrated that elevated cytoskeletal tension in KDFs promotes lysosomal exocytosis. Taken together, these findings provide new mechanistic insights into the altered mechano-sensing behaviour of KDFs and identify a novel biomechanical switch that drives lysosomal remodelling.

Consistent with our findings, previous studies have reported similar increases in cytoskeletal tension and an upregulation of YAP/TAZ signalling, a key mechanotransduction component of the Hippo pathway, in keloid fibroblasts (Aramaki-Hattori et al. 2017; Gao et al. 2021). Here, we provide new evidence for a functional linkage to dysregulated ECM deposition and exocytic pathways. Constitutively high levels of cytoskeletal tension in KDFs may therefore contribute to excess ECM production within keloid scars. However, an important unanswered question is how and when KDFs acquire this heightened mechanical response. Several recent reports have described the ability of cells to retain a mechanical memory of previous environments (Li et al. 2017; Yang et al. 2014), suggesting that the stiffened scar ECM in vivo could play a role in establishing the mechanical phenotype of KDFs. Nevertheless, the cells used in our studies were all expanded for several passages on rigid tissue culture plastic, indicating that the mechanical phenotypes of KDFs are deeply encoded and may reflect additional genetic or epigenetic mechanisms. In future studies, it will be interesting to explore the responses of fibroblasts from keloids and adjacent normal skin of the same patient, as well as the response of KDFs and NDFs prior to expansion on plastic surfaces.

A major challenge in keloid research is the high level of inter-patient heterogeneity, which was evident in the RNA-seq analysis conducted in this study. Here, we observed strong clustering of transcriptional profiles according to specific donors, and standard protocols (e.g. DESeq2) failed to identify DEGs regulated by substrate stiffness. Instead, we used the higher PCs (PC5) to uncover the mechanically regulated genes and pathways, and this less conventional approach successfully identified both classic mechanotransduction pathways and novel pathways, such as autophagy. It will be interesting in future studies to explore some of the additional mechanically regulated pathways, such as Rap1 and cytokine signalling. In addition, it will be important to expand the transcriptional analysis to a greater number of keloid patients and investigate potential correlations with body site, sex, and ethnicity, which were not accounted for in this study due to the limited sample size.

To the best of our knowledge, the biomechanical regulation of exocytosis has not been previously implicated in keloid scar formation. Excessive lysosomal exocytosis has been linked to increased ECM and matrix metallo-proteinase secretion in other forms of fibrosis, including patients with sialidosis and pulmonary fibrosis, and in mice lacking the negative regulator of lysosomal exocytosis, *Neu1* (van de Vlekkert et al. 2019). In addition, intracellular transport of lysosomal vesicles and fusion with the plasma membrane depend on F-actin and myosin activity (van Bommel et al. 2019; Encarnação et al. 2016), and microtubules and the actin cytoskeleton facilitate lysosomal remodelling in establishment of leader cells in migrating epithelia (Marwaha et al. 2023). It is therefore possible that elevated cytoskeletal tension in KDFs could contribute to scar formation via increased lysosomal exocytosis and release of key scar components or regulatory factors. For example, IL-6 secretion is upregulated in keloid fibroblasts and drives supracellular alignment of the actin cytoskeleton and ECM (Kenny et al. 2023), but the relationship to lysosomal remodelling has yet to be determined. Further insights into the specific secreted factors and regulatory mechanisms involved in these processes could therefore lead to new therapeutic targets and treatments for controlling keloid scar formation.

## Materials and methods

### Cell source and culture

Primary dermal fibroblasts were previously isolated from redundant skin of normal healthy donors or excised keloid scars. Donor samples K1, K3, K5, K6, N1 and N3 were obtained from keloid patients and healthy volunteers from the plastic surgery department at Barts Health NHS Trust. All subjects gave informed consent, and the study was conducted under local ethical committee approval (East London Research Ethics Committee, study 2011-000626-29). Patient samples K2, K4 and N2 were obtained from the St. George’s NHS Healthcare Trust (North of Scotland REC, study 14/NS/1073). Available patient information is listed in below (Table 1).

**Table 1:** Information of keloid patients and healthy skin donors.

| Sample | Sex | D.O.B | Body location | Ethnicity |
| --- | --- | --- | --- | --- |
| K1 | Female | Unknown | Ear | Afro-Caribbean |
| K2 | Female | 2002 | Ear | Unknown |
| K3 | Male | Unknown | Unknown | Afro-Caribbean |
| K4 | Female | 1983 | Earlobe | Unknown |
| K5 | Unknown | Unknown | Ear | Afro-Caribbean |
| K6 | Male | Unknown | Cheek | Afro-Caribbean |
| N1 | Female | Unknown | Breast | Caucasian |
| N2 | Female | Unknown | Abdomen | Caucasian |
| N3 | Male | Unknown | Scalp | Unknown |

All materials and reagents were from Thermo Fisher Scientific unless otherwise stated. Cells were cultured in Dulbecco’s Modified Eagle’s Serum (DMEM) with GlutaMax supplemented with 10% Fetal Bovine Serum (FBS) (Biosera) and 1% penicillin/streptomycin (P/S) at 37°C with 5% CO_2_. Medium was changed every 2/3 days and passaged once they reached confluency. Primary fibroblasts were used for experiments between passages five and nine.

### PA gel fabrication

Large rectangular (25×60 mm) coverslips and small circular (13 mm) coverslips were cleaned using the Henniker plasma HPT-200. Both sides of the coverslips were exposed to the plasma for 10 minutes. PA gels were prepared, and stiffness was measured as previously described in Laly et al., 2021.

### Cell derived matrices

Small PA hydrogels were seeded with 1×10^5^ cells and incubated for 5 days with the addition of 50 μg/ml ascorbic acid each day. On day 5 the cells were washed with 1X phosphate-buffered saline (PBS) and then 1 ml of pre-warmed extraction buffer (1X PBS, 0.25% triton X-100 and 10 mM NH_4_OH) was added to each coverslip and left for four minutes to allow the cells to lyse. After four minutes half of the extraction buffer was removed and replaced with 1X PBS and this step was repeated until no cells were visible. The DNA was digested using 10 μg/ml DNase 1 (Roche), incubated for 30 minutes at 37°C, 5% CO_2_. Matrices were then washed two more times with 1X PBS and were stored at 4°C in PBS with P/S prior to immunofluorescent staining.

### Western blotting

Large PA hydrogels or six-well-plates were seeded with 3×10^5^-4×10^5^ cells per gel/well and incubated for 24 hours. Cells were washed with 1X PBS and cell lysis was performed on ice using 75 μl of radio immunoprecipitation assay (RIPA) buffer, containing 1X PhosSTOP (Roche) and 1X protease inhibitor (Sigma Aldrich) per gel/well. Cell scrapers were used to help detached and lyse the cells. Lysates were sonicated for 30 seconds on/off for 10 minutes and centrifuged at 4°C, 12500 rpm for 10 minutes. The supernatant for each lysate was collected ready for western blotting analysis. Each sample was then diluted with ddH_2_O and 4X NuPAGE sample buffer (containing 1% β-mercaptaethanol). Each protein sample was heated to 80°C for five minutes.

Equal amounts of protein were loaded onto precast 4-15% gradient SDS-PAGE gels (Bio-Rad) and run at 100V for between 1 – 1 ½ hours. Gel transfers were either performed at 300A for one hour onto a 0.45 μm (0.2μm for LC3 blots) nitrocellulose membrane (GE Healthcare) or transferred using the Trans turbo transfer system (Bio-Rad). Blots were blocked with 5% milk in Tris-buffered saline with 0.1% Tween (TBS-T) or 5% BSA in TBS-T according to the primary antibody’s diluent. Primary antibodies were applied overnight at 4°C, rotating. Blots were then washed with TBS-T three times for five minutes and incubated at room temperature with secondary antibodies for one hour, rotating. Blots were then washed with TBS-T three times for five minutes and exposed to Immobilon Western Chemiluminescent HRP substrate (Merck Millipore) according to product instructions for 30 seconds before being developed. Primary antibodies included anti-GAPDH (clone 124H6, Cell Signalling Technology [1:20000]), anti-LC3B (Cell Signalling Technology [1:200]) and anti-SQSTM1 (p62, Novus BioH00008878-M01 [1:500]).

### RNA-sequencing

Large coverslip PA hydrogels were seeded with 4×10^5^ cells from three different normal and keloid patient samples and cultured for 24 hours. Medium was then removed from each gel and washed with 1X PBS while being transferred over to new 4-well plates. Using RNeasy® plus micro kit (Qiagen), 350 μl of RLT lysis buffer was added directly onto the PA hydrogels and washed over three times. The lysates were collected, and the RNA was extracted and purified as per manufacturer instructions. Samples were quantified using the NanoDrop™ 1000, and QC assessment, library preparation and sequencing were performed by Lexogen GmbH. The QuantSeq 3’ mRNA-seq Library Prep Kit FWD (Lexogen) was used to generate Illumina compatible library sequences, and sequencing was performed with the Illumina NextSeq high output sequencing system.

Sequencing data in FastQ files were aligned to the human genome (annotation hg19) HISAT2 on the Galaxy platform. Differential expression (DE) analysis was conducted using the graphic user interface SeqMonk (https://www.bioinformatics.babraham.ac.uk/projects/seqmonk/). The data were quantified using the RNA-seq quantitation pipeline and DE analysis was performed between the four conditions (NDF 8kPa, NDF 214kPa, KDF 8kPa and KDF 214kPa). Principle component analysis (PCA) was conducted on the differentially expressed genes (DEGs) and the highest weighted PC5 probes were extracted by filtering based on their absolute rotation value (≥10). Gene set enrichment analysis (GSEA) was performed using Webgestalt (Liao et al. 2019). Default settings were used except the number of categories generated, which was changed to 20. The protein-protein interaction network was produced using the STRING database (Szklarczyk et al. 2023). A full STRING network was produced meaning the edges indicate both functional and physical protein associations, in addition the edges represent confidence in the interactions whereby the thicker the line indicates the strength of data support. Raw and processed RNA-seq data was deposited in the Gene Expression Omnibus (GSE 246562).

### Immunostaining

Small PA gels were seeded with 1×10^4^-2×10^4^ cells and incubated for 24 hours. Cells were fixed with 4% paraformaldehyde (PFA, Sigma Aldrich) for 10 minutes and permeabilised with 0.2% Triton (Sigma Aldrich) in 1X PBS for five minutes. Cells were incubated for 30 minutes at room temperature with blocking solution (10% FBS in 1X PBS). Primary antibody staining was performed at 4°C, overnight. Cells were then washed three times with 1X PBS followed by a one-hour incubation with secondary antibodies/dyes solution. Cells were washed three times with 1X PBS and once with ddH_2_O. To prepare PA samples for imaging, 10 μl Mowiol (Sigma) was pipetted onto small square (22×22mm) coverslips and the PA gels were mounted on top (PA gel/cells facing down). Mounted samples were left to dry in the dark and once dry the edges were sealed with clear nail polish. 10 μl Mowiol was added to microscope slides and the small square coverslips with the PA gels were mounted on top (PA gel in the middle) and left to dry overnight in the dark. To prepare non-PA gel samples for imaging 10 μl Mowiol was pipetted onto a microscope slide and samples were mounted on top (cells facing down). Normal skin and keloid tissue samples were fixed with 4% paraformaldehyde, dehydrated, and embedded in paraffin, and tissue sections were cut to 7 μm immediately before immunostaining. Paraffin sections were dewaxed, and heat-mediated antigen retrieval was performed with 10mM sodium citrate. Tissue sections were blocked and stained with primary and secondary antibodies as described above. Primary antibodies included anti-paxillin (clone 177, BD Biosciences [1:200]), anti-fibronectin (Sigma F3648 [1:1000]), anti-collagen (Abcam ab6308 [1:1000]), anti-LAMP1 (Cell Signaling 9091S [1:100]), anti-LAMP2 (Santa Cruz H4B4 [1:100]), and anti-LAMP1-LD (R&D Systems AF4800 [1:100]). Secondary antibodies and dyes included phallodin Alexafluor 488 (Molecular Probes [1:1000]), phalloidin Alexafluor 555 (Molecular Probes [1:1000]), anti-mouse Alexafluor 555 (Molecular Probes [1:1000]), anti-rabbit Alexafluor 488 (Molecular Probes [1:1000]), and anti-sheep Alexafluor 488 (Molecular Probes [1:1000]).

### Fluorescence microscopy and image analysis

F-actin/FA and CDM immunofluorescence images were acquired using Zeis 880 Laser Scanning Confocal at 63X magnification. Fast Airyscan was used to capture Z-stacks which were processed into maximum intensity projections. For each condition, 10 fields of view (FoV) were taken. Cell mask, LAMP1, and F-actin (after inhibitor treatment) immunofluorescence images were acquired using Leica DM5000B Epi-Fluorescence microscope. When imaging cell mask, 10 FoVs were taken per condition at 10X magnification, and F-actin and LAMP1 were imaged at 63X magnification.

The IN Cell Developer Toolbox (GE) was used to analyse the F-actin filaments and FAs with specific segmentation kernal sizes (F-actin filaments = 5, FAs = 7) and acceptance criteria (F-actin filaments: density levels >600 and maximum chord curve >20, FAs: density levels >800 and maximum chord curve >25 but <250). To calculate cell area, the F-actin and FA channels were overlaid to act as an outline of the cells. When analysing cell area and aspect ratio based on cell-mask staining using the IN Cell Developer Toolbox Software, each cell was detected based upon the presence of a nuclei and the intensity of the cell mask stain (segmentation set to intensity, minimum threshold = 22 pixels, maximum threshold 255 pixels).

ImageJ was used to analyse CDMs and LAMP1. The local thickness tool was utilised to determine fibre spacing whereby the binary images were inverted. Total fluorescent area and fluorescence intensity were also calculated by ImageJ. To calculate the perinuclear LAMP1, the nuclear DAPI image was used as a mask and dilated 50%. The percentage of total LAMP1 fluorescence per cell within the dilated nuclear mask was classified as perinuclear. For quantification of exocytic vesicles, the particle analysis tools within ImageJ was used to calculate the number of LAMP1-LD particles per cell.

### Statistical analysis

All statistical analysis was performed using GraphPad Prism. Results were averaged per experiment then averaged again per biological or independent experimental repeat. Acquired data were subjected to either a two-way analysis of variance (ANOVA) or an unpaired t-test, with significance set at p-value ≤0.05.

## Data Availability Statement

RNA-seq data is available through the Gene Expression Omnibus (Accession number GSE 246562). All other raw or processed data will be made available upon request.

## ORCiDs

RR: 0009-0008-8248-0752

DB: 0000-0002-3955-1091

TJS: 0000-0001-6187-368X

JTC: 0000-0002-5955-8848

## Acknowledgements

This work was funded by the Barts Charity (Grant G-002184) and PhD studentship from the Biotechnology and Biosciences Research Council (London Interdisciplinary Doctorate programme).

## Author Contributions

Investigation: RR and YG. Formal Analysis: RR, RNW, JTC. Methodology: DB. Resources: DB and TJS. Writing – original draft: RR and JTC. Writing – review and editing: RR, TJS, DB, JTC. Conceptualization: TJS and JTC. Supervision: JTC. Funding Acquisition: JTC.

## Supplementary Information

**Figure S1:**
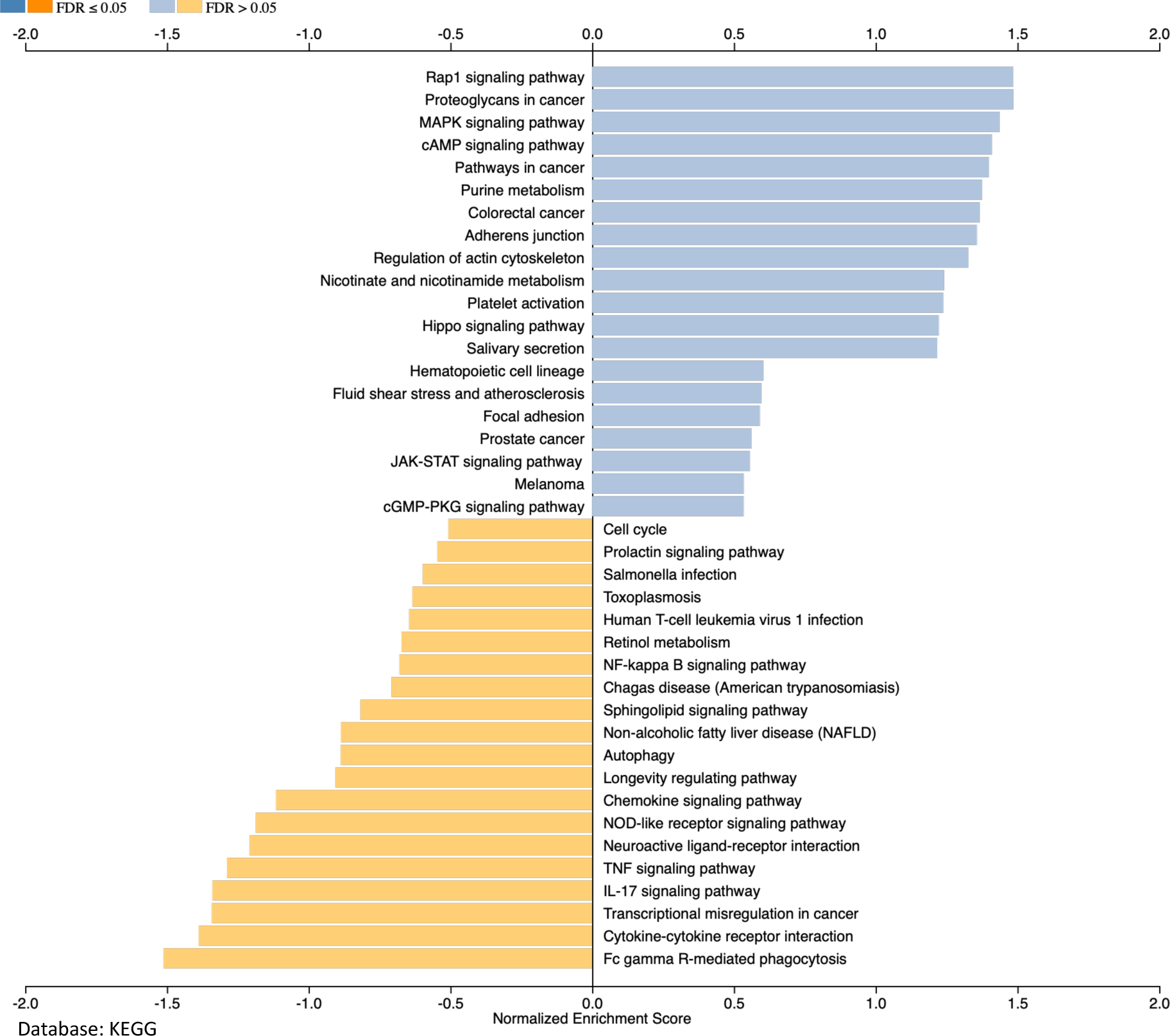
Gene set enrichment analysis (GSEA) of differentially expressed genes from PC5 with absolute rotation >10. Analysis was performed using the SeqMonk platform and KEGG database, and data represent the normalised enrichment score for each pathway.

**Figure S3:**
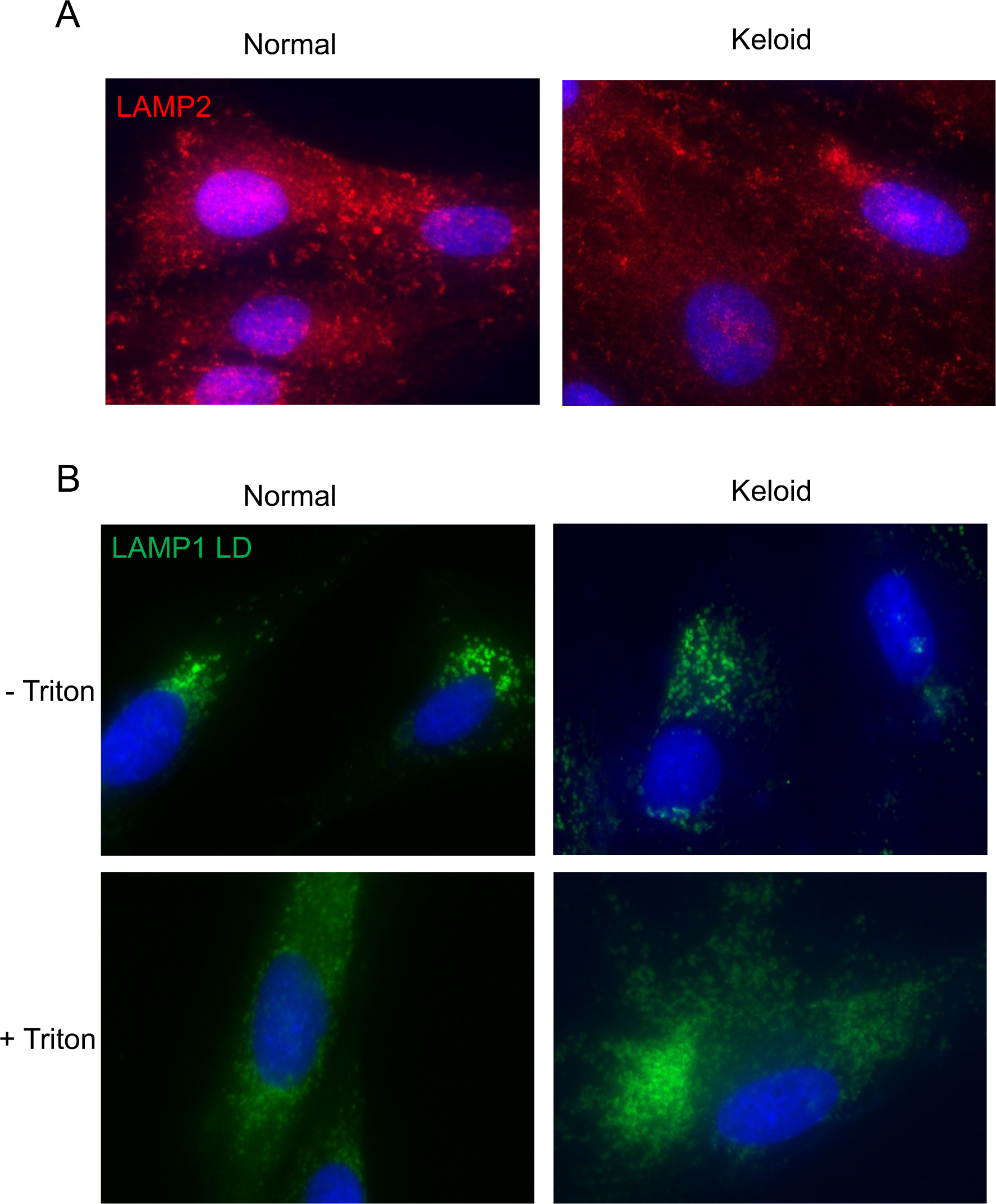
(A) Representative images of NDFs and KDFs stained for LAMP2. (B) Representative images of NDFs and KDFs stained for LAMP1-LD with or without permeabilisation (0.2% Triton-X100 for five minutes).

